# Individual Brain Structure Deviations and its Gene Expression Signatures in Early-Onset Schizophrenia

**DOI:** 10.64898/2026.02.06.704304

**Authors:** Yun-Shuang Fan, Yuting Xu, Yong Xu, Liju Liu, Mi Yang, Jing Guo, Huafu Chen

## Abstract

**Background:** Schizophrenia is a highly heritable mental disorder associated with widespread anatomical alterations during neurodevelopment. Converging evidence suggests transcriptomic architecture underlying brain abnormalities in schizophrenia, while how individualized brain morphological deviations relate to gene expression levels remains unclear.

**Methods:** To investigate individual-level brain deviations and its transcriptomic signatures in schizophrenia, this study collected T1-weighted MRI data from 95 early-onset schizophrenia (EOS) patients and 99 typically developing (TD) controls. Normative modeling was used to measure individual deviations in cortical thickness and subcortical volume. Partial least squares regression was calculated to capture covarying patterns between structural deviations and whole-brain gene expression levels. Clustering analysis was performed on latent brain-gene covarying components, and the results were further functionally decoded through gene enrichment analyses.

**Results:** Group-level comparisons suggested patients with EOS showed consistently decreased z-scores of cortical thickness in the frontal and temporal lobe regions, while increased inter-individual variability in the lingual gyrus. Clustering analysis of z-scores with transcriptomic signatures identified two distinct brain-gene covarying subtypes. Subtype 1 showed thickening cingulate gyrus, thinning occipital pole, and atrophic subcortical nuclei. Subtype 2 exhibited widespread cortical thinning across the frontal, parietal, temporal, and limbic regions, but enlarged subcortical nuclei. Genes underlying two subtypes were both enriched for neurodevelopmental diseases. However, subtype 1 was associated with synaptic transmission, and subtype 2 was related to cytoskeletal and neuronal connectivity.

**Conclusion:** This study reveals individual-level anatomical deviations and transcriptomic heterogeneity in early-onset schizophrenia. The findings provide an individualized brain-gene coupling framework for understanding pathophysiology of schizophrenia during brain development.

## Introduction

Schizophrenia is a polygenetic psychiatric disorder characterized by extensive cortical thinning predominantly in the frontal and temporal regions and enlarged bilateral lateral ventricles^1–4^. These brain structural abnormalities in schizophrenia have been reported to relate to disrupted synaptic remodeling during brain developement^5–7^. Early-onset schizophrenia (EOS), which was neurobiologically continuous with its adult counterpart, offers a chance to reveal the interface between pathophysiologic and neurodevelopmental progresses. However, adolescence is also a period of excessive cortical pruning and insufficient subcortical pruning, according to schizophrenic hypotheses, including the synaptic hypothesis, during which the brains of adolescents exhibit greater inter-individual variability than those of adults^6,8–10^.

Although strong evidence reveals group-level differences of brain structure in EOS, it is increasing suggested that these differences vary across individuals^11–14^. Inter-individual variability in brain structural abnormalities across EOS patients has been reported in brain structural abnormalities, including cortical thickness and subcortical volume alterations^15–17^. The meta-analysis study indicated that patients with schizophrenia exhibited increased consistency in anterior cingulate cortex volume, while greater variability was observed in putamen, temporal lobe, thalamus and third ventricle volume^9^. Another study also revealed higher heterogeneity in schizophrenia regarding cortical thickness and area, cortical and ventricle volume, and hippocampal subfield^8^. Therefore, while group-level comparisons have established a robust pattern of brain structural alterations in schizophrenia, their reliance on population averages may obscure the substantial variability that exists across individuals. This limitation underscores the need for capturing and quantifying individual-level deviations from normative brain development.

Recently, the normative modeling approach has been proposed to quantify individual-level neurobiological deviations in brain disorders, such as major depressive disorder, autism spectrum disorder and Alzheimer disease^18–20^. After establishing a normative model based on the healthy population, it is feasible to measure individual-level structural deviations from the normative range. For example, the hippocampus has been identified as the region exhibiting the highest heterogeneity among patients with Alzheimer’s disease^21^. Likewise, transdiagnostic brain abnormalities were found between individuals with schizophrenia and bipolar disorder in a few areas, e.g., the frontal, temporal, and cerebellar regions using normative model^22^. In schizophrenia, widespread negative deviations have been found in fractional anisotropy and cortical thickness, affecting almost all white matter tracts and cortical regions^23^. Accordingly, the normative modeling approach could be used to measure how EOS subject diverges from typical brain development and their inter-individual variability.

Notably, recent advancement in transcriptomics has enabled us to integrate multimodal neuroimaging and genomics data^24–27^. Specific sets of genes have been identified as playing crucial roles in disease susceptibility,and these genes are also involved in synaptic plasticity, neurodevelopment, and immune-inflammatory responses^28–30^. The relationship between transcriptomic profiles and brain alterations has been reported in multiple brain disorders^31^. For example, brain functional alterations of major depressive disorder have been shown to be associated with transcriptomes of a set of genes enriched in transsynaptic signaling and calcium ion binding^32^. Gene expression levels of neurodevelopmental disease risk genes have been shown to be related with brain structural alterations in EOS^33^. However, previous studies were based on group-level alteration patterns, obscuring the molecular biological mechanisms underlying individual-level brain deviations.

To elucidate individualized anatomical deviations and their genetic relevance in schizophrenia during brain development, this study utilized a cohort of EOS patients (defined by onset age before 18 years), along with whole-brain gene expression maps from the Allen Human Brain Atlas microarray dataset (https://human.brain-map.org/). We first employed partial least squares (PLS) regression to associate individual-level deviation patterns from normative model in cortical thickness and subcortical volumes with whole-brain gene expression profiles, thereby deriving subject-specific PLS components that reflect brain–gene coupling characteristics. Clustering analysis was then performed on PLS components across individuals to identify EOS subtypes that differed in both brain structural and molecular characteristics. To further reveal the biological mechanisms underlying these subtypes, we conducted disease enrichment analysis, Gene Ontology (GO) enrichment analysis, and Kyoto Encyclopedia of Genes and Genomes (KEGG) pathway enrichment analysis. We hypothesized that EOS patients exhibit biological subtypes with distinct transcriptomics and neurobiological processes under the inter-individual variabilities in brain structural changes.

## Methods

### Participants

A total of 194 participants including 95 antipsychotic-naive first-episode EOS patients and 99 typically developing (TD) controls were recruited from the First Hospital of Shanxi Medical University and the local community through advertisements, respectively. All participants were aged between 7 and 17 years and were right-handed. The schizophrenia diagnosis followed the criteria in the Structured Clinical Interview for DSM-IV and were confirmed by at least one senior psychologist after a minimum 6-month follow-up period. The psychiatric symptomatology of 71 EOS patients was assessed using the Positive and Negative Syndrome Scale (PANSS). Exclusion criteria for all subjects included neurological MRI anomalies, electronic/metal implants, or histories of substance abuse. EOS patients were also excluded if they had been suffering from the illness for more than 1 year. TD controls were excluded if they had a family history of psychiatric disorders. This study was approved by the Ethics Committee of the First Hospital of Shanxi Medical University. Written consent was obtained from every participant and their parents or legal guardians.

### Imaging data acquisition

T1-weighted MRI data were acquired using a 3 Tesla MRI scanner (MAGNETOM; Siemens, Germany) using a three-dimensional fast-spoiled gradient echo sequence with the following parameters: repetition time (TR) = 2300 ms; echo time (TE) = 2.95 ms; flip angle = 90°; matrix = 256 × 240; slice thickness = 1.2 mm (no gap); and voxel size = 0.9375 × 0.9375 × 1.2 mm^3^; and 160 axial slices.

### Imaging data processing

T1-weighted structural data were preprocessed using FreeSurfer (v7.1.0, https://surfer.nmr.mgh.harvard.edu/), which involved cortical segmentation and surface reconstruction. The preprocessed images were then registered to the MNI152 template and mapped onto the Destrieux cortical atlas using the Ciftify package (v2.3.3) [30]^34^. The subcortical nuclei were segmented by using FreeSurfer automatic subcortical segmentation atlas^35^. For subsequent analysis, we selected 16 subcortical areas for subsequent analysis, including the bilateral accumbens area, amygdala, caudate, hippocampus, pallidum, putamen, thalamus, and lateral ventricle.

### Normative Modeling

To capture EOS individual regional deviation with the covariates including age and sex, we built normative models for our data. To obtain a normative model adapted for our location, we split the sample of 99 TD controls into an adapting set (N = 40) and a testing set (N = 59), which were matched for sex and age. As previously suggested, our sample size of adapting cohort, which was larger than five, ensured a reliable representative distribution of participants for our center^36–38^. Additionally, we used another testing dataset including 95 EOS patients for the same center. We first trained region-wise normative models (i.e., one model per region) for cortical thickness and subcortical nuclei grey matter using the adapting dataset and Bayesian linear regression via the Predictive Clinical Neuroscience toolkit package (PCNToolkit, version 0.33.0)^36^. The resulting models were used to evaluate region-wise deviation maps for each participant in the testing dataset relative to the normative population (Figure 1A). Specifically, for brain region d (cortical thickness or subcortical volume), the deviation for subject n was quantified by the Z_nd_ score^39^:

**Figure 1.**
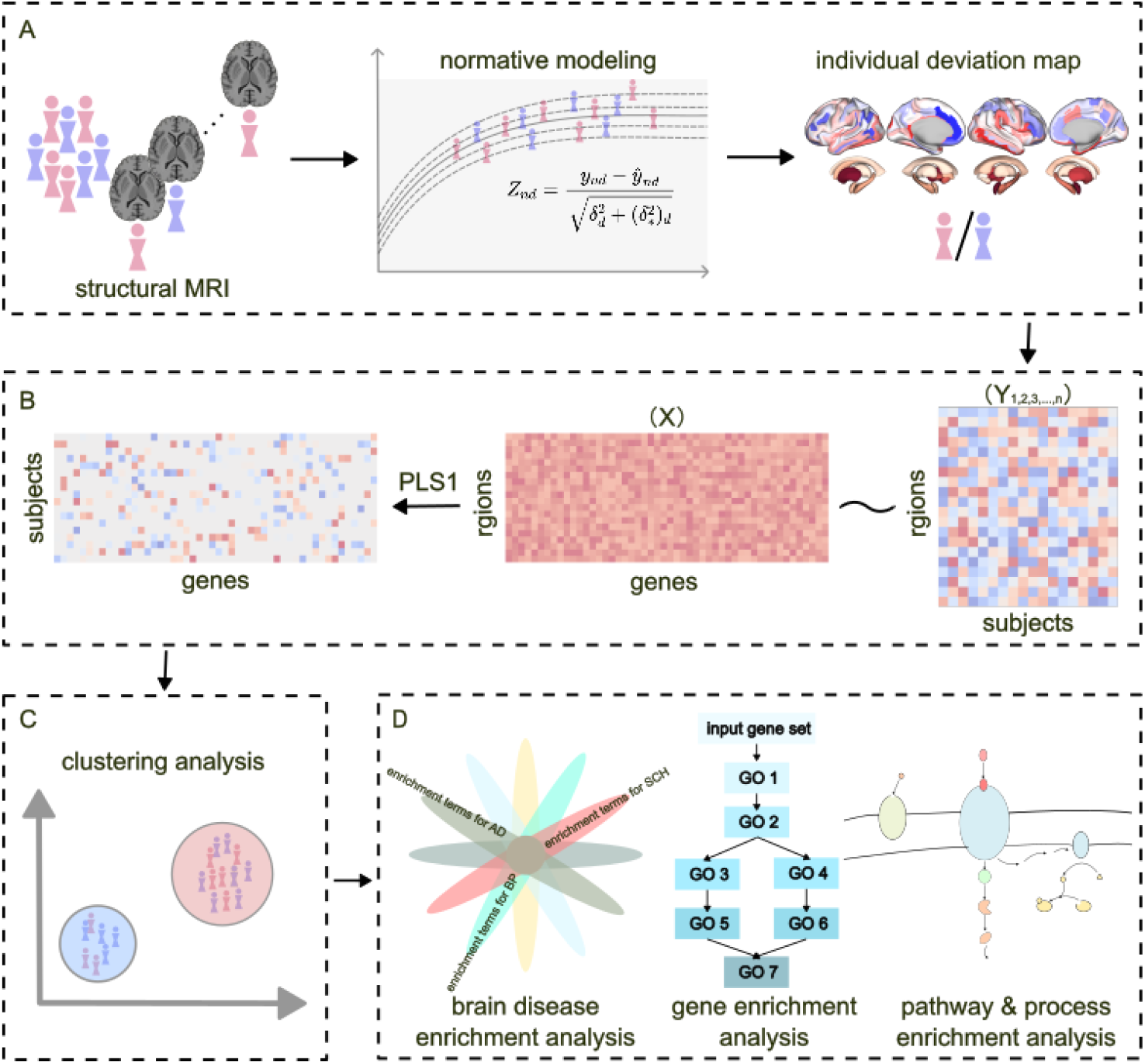
Flowchart of the proposed method. **(A)** Schematic overview of the methodology combining structural MRI with normative modeling to generate individual deviation maps. **(B)** PLS analysis framework showing the correlation between structural deviations and gene expression. **(C)** Clustering analysis based on PLS1 results, grouping subjects into distinct clusters based on their gene expression profiles. **(D)** Overview of the gene enrichment analysis pipeline. Input gene sets are analyzed for enrichment across various biological categories, including brain disease associations (e.g., Alzheimer’s Disease, Schizophrenia), Gene Ontology, and Kyoto Encyclopedia of Genes and Genomes pathway. Structural MRI, structural magnetic resonance imaging; PLS, partial least square; PLS1, the first partial least square component.

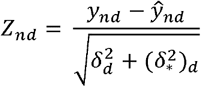

Where *y*_*nd*_ and *ŷ*_*nd*_ are observed values and model-predicted values for brain features, respectively. 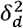 is the estimated noise variance reflecting uncertainty in the data, and 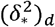 is the variance attributed to modeling uncertainty. Finally, for n subjects in the EOS group, we obtained the *Z-scores* vector Y (169 brain regions × 1) that contains individual regional deviation measurement.

### Partial Least Squares Regression

The Allen Human Brain Atlas dataset (https://human.brain-map.org/) provides access to microarray transcriptomic data from six postmortem adult human brains without history of psychiatric disease^40^. The regional gene expression levels were preprocessed down-sampled to the Destrieux atlas with subcortical nuclei using the pipeline provided by the *abagen* toolbox^41^. We obtained a gene expression matrix (169 brain regions ×15,633 genes).

To access individual anatomical deviation-related gene profiles, we calculated PLS regression between the gene expression matrix and *Z-score* deviation matrix for each EOS patient^42^. PLS regression is a multivariate method that jointly decomposes two matrices, here X (169 × 15,633 gene expressions) and Y (169 × 1 cortical thickness or subcortical volume *Z-score*s), into latent components. The first PLS component (PLS1) is the linear combination of gene expression map most strongly correlated with individual *Z-score map* across 169 regions (**Figure 1B**). The statistical significance of the variance explained by PLS1 was assessed using a bootstrapping approach (5000 iterations)^43^. In each individualized PLS regression, significant genes were ranked based on their contribution to PLS1^44^. Specifically, the gene weight vectors were derived from the PLS1 analysis, where the Z-value for each gene was calculated to quantify its contribution to the individual deviations across EOS patients, The Z-value represents the standardized weight, reflecting the gene’s relative importance in explaining the variance in neuroanatomical features. These gene weight vectors were then merged into a two-dimensional subject × gene weight matrix.

### Clustering analysis

To identify transcriptomics-related subtypes of EOS, we performed clustering analysis on the two-dimensional subject × gene weight matrix. We first performed quality control on the original two-dimensional “subject × gene” gene weight matrix. Specifically, since high-dimensional gene expression data is inherently sparse, an excess of zero expression values can significantly affect the stability of distance calculations and the reliability of the clustering structure. To reduce the high proportion of zero values caused by sparse expression and technical dropout, we excluded subjects with 0 PLS1 gene expression. For the filtered matrix, since the feature dimension is still extremely high, directly clustering it may lead to distance metric degradation and the “curse of dimensionality” problem. Therefore, we used truncated singular value decomposition (truncated SVD) to linearly reduce the dimensionality of the gene expression matrix, extracting the main variable components and reducing the noise dimension^45,46^.

After dimensionality reduction, to further improve the clustering performance on highly sparse data, we performed L2 normalization on each truncated SVD representation, ensuring that all sample vectors are distributed on a unit sphere. At this point, Euclidean distance and cosine similarity maintain consistent ranking, thus K-Means clustering analysis can be considered an approximate spherical clustering method, better suited for capturing directional differences in gene expression. To determine the optimal dimensionality reduction dimension and number of clusters, we iterate through the SVD dimension range from 2 to 20, evaluating the spherical K-Means clustering performance for each dimension with cluster numbers k ranging from 2 to 9, using the silhouette score as the clustering quality metric^47^. For each truncated SVD dimension, we select its corresponding optimal number of clusters and highest score, ultimately determining the globally optimal dimensionality reduction dimension and number of clusters through cross-dimensional comparison. Based on this optimal combination, we perform a final spherical K-Means clustering analysis to obtain the cluster labels for the subjects. To illustrate the clustering structure and help explain the inter-cluster separation, we additionally used uniform manifold approximation and projection (UMAP), t-distributed stochastic neighbor embedding (t-SNE), and the first two dimensions of truncated SVD projection for two-dimensional visualization (**Figure 1C**).

### Gene enrichment analysis

To explore the molecular and cellular mechanisms involved in transcriptomics-related subtypes, we performed functional annotation and pathway enrichment analyses on the characteristic gene sets of each subtype (**Figure 1D**). For each subtype, we first calculated the frequency of gene occurrence (i.e., the proportion of subjects in a specific subtype where the gene has a non-zero Z-value, based on its significant association in the PLS1 analysis), and selected genes with an occurrence rate greater than 15% for subsequent enrichment analysis, aiming to reduce noise from sparsely expressed genes^48^.

Subsequently, gene function enrichment analysis was performed using the Metascape platform (https://metascape.org/gp/index.html#/main/step1) platform. The gene lists derived from each subtype were used for over-representation analysis of brain diseases, GO and KEGG pathway enrichment. In the GO enrichment portion, we simultaneously evaluated three ontologies—Biological Process, Molecular Function, and Cellular Component-identify subtype-specific biological processes, molecular functions and cellular structures. KEGG pathway analysis was performed to identify relevant signaling pathways and cellular processes. Metascape used hypergeometric tests to assess enrichment significance, with the false discovery rate (FDR) controlled by the Benjamini-Hochberg procedure. Significantly enriched terms were identified based on the criteria of p < 0.01, an enrichment factor greater than 1.5, and at least three genes contributing to the enrichment.

Enrichment results were visualized using multiple approaches, including bubble plots for brain diseases and GO terms, showing gene proportions and statistical significance, KEGG pathway clustering networks illustrating functional relationships among pathways, and interactive gene-function mapping networks. These complementary analyses systematically delineate the neurobiological pathways and cellular processes potentially involved in the different expression subtypes.

## Results

### Brain structural deviations in EOS

To measure EOS deviations in cortical thickness and subcortical volumes from normative data, we applied regional normative models. Patients with EOS showed individual-specific positive or negative deviation *Z-score* distributions across brain regions (**Figure 2A**). After averaging the deviation maps within the EOS group, we identified a group-level EOS pattern, with predominantly negative deviations in cortical thickness and positive deviations in subcortical volumes. We then performed two-sample *t*-tests between the *Z-score* maps of patients and controls, and observed significantly reduced *Z-score* (q_FDR_ < 0.05) in the temporal, frontal and parietal lobes.(See **Table S1** for detailed statistical values). In terms of subcortical nuclei, patients only showed increased *Z-score* in the left lateral ventricle (*t-value* = 2.87, *q*_*FDR*_ = 0.03) (**Figure 2B**). Based on the results of the standard deviation comparison between the HC and SZ groups, significant differences in brain regions were observed in the left hemisphere’s medial occipito-temporal gyrus in which it indicated that the heterogeneity in the SZ group was predominantly concentrated.

**Figure 2.**
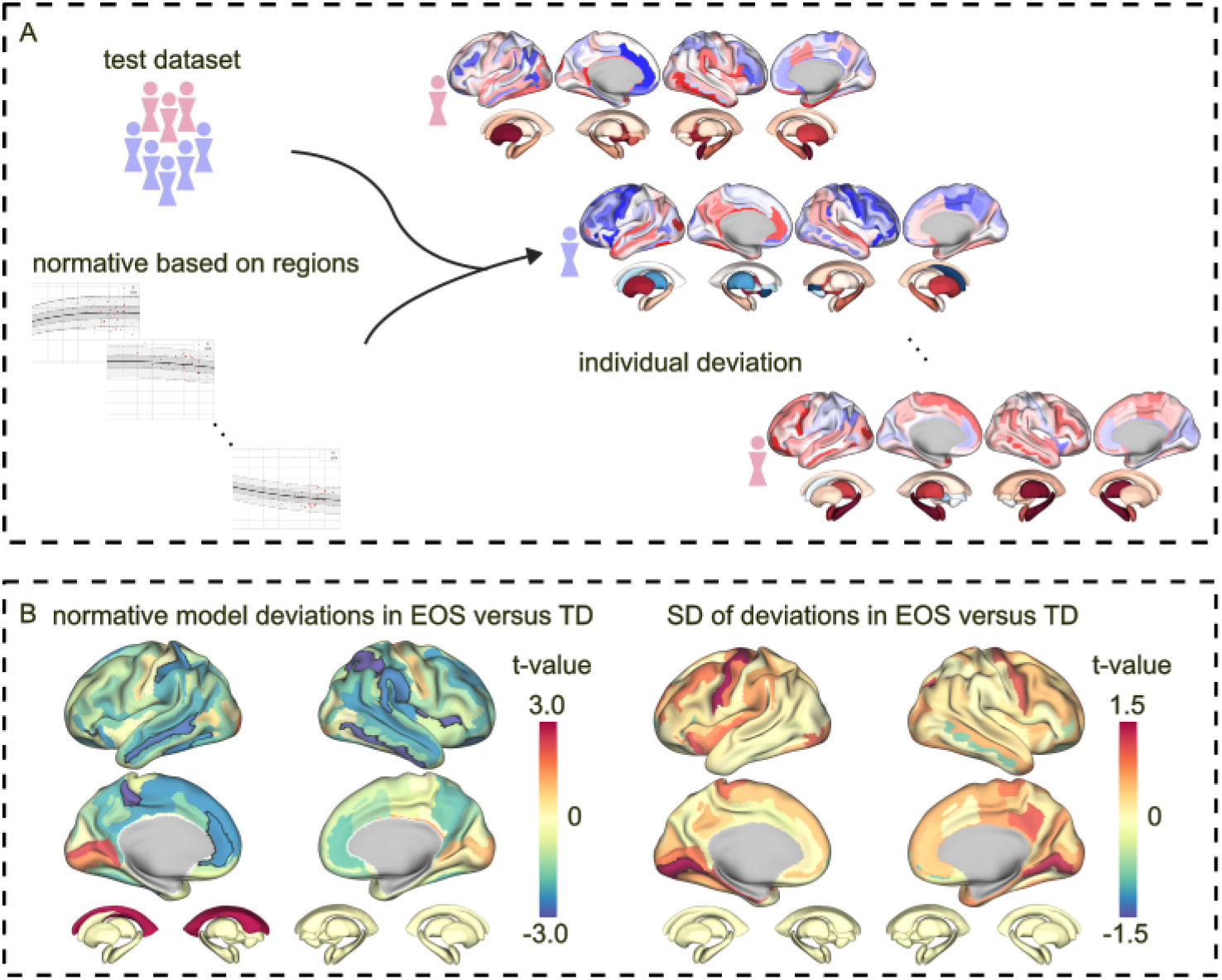
Brain structural deviation patterns in EOS patients. **(A)** Overview of the normative modeling approach used to assess structural deviations in EOS patients. The model compares individual brain regions from the test dataset to normative data, generating personalized deviation maps. **(B)** (*Left*) Group-level comparison between patients and TD controls. The map shows t-values across the entire brain, with regions highlighted in red/blue indicating significant positive/negative deviations, respectively. Outlines in black demarcate brain regions that survived FDR correction *(P < 0*.*05)*. (*Right*) Group comparison of regional deviation variability (standard deviation, SD) between EOS patients and TD controls, with black outlines similarly indicating regions that remained significant after FDR correction. These patterns reveal spatially specific structural alterations in EOS. EOS, early-onset schizophrenia; TD, typically developing.

### Transcriptomics-based EOS subtype groups

To investigate transcriptomic association of EOS deviation maps, we conducted individual-level PLS analysis on structural deviation and gene expression matrices. We identified individual-specific significant gene sets underlying the individual structural deviation map (**Figure 3A**). After clustering gene loading maps across patients, we found that the optimal number of clusters was 2, with the corresponding maximum silhouette coefficient was 0.721, suggesting the reliability of clustering^47,49^. Subtype 1 showed dominantly positive cortical deviations with atrophy of subcortical nuclei, while subtype 2 showed widespread cortical thinning and enlargement of subcortical nuclei (**Figure 3B**). However, after FDR correction, subtype 1 only showed significant negative deviation effects in the left posterior-ventral part of the cingulate gyrus (*t-value* = -3.55, *q*_*FDR*_ = 0.01) and the left hippocampus (*t-value* = -2.84, *q*_*FDR*_ = 0.04), right hippocampus (*t-value* = -3.84, *q*_*FDR*_ = 0.007), left thalamus (*t-value* = -3.72, *q*_*FDR*_ = 0.01), right thalamus (*t-value* = -3.99, *q*_*FDR*_ = 0.0067), left amygdala *(t-value* = -2.84, *q*_*FDR*_ = 0.04). In contrast, subtype 2 exhibited significantly negative deviation effects on the frontal, parietal, and temporal lobes, with minor involvement of the occipital lobe, cingulate and limbic systems, insula, and orbitofrontal region (See **Table S2 & S3** for detailed statistical values). Moreover, patients in subtype 2 showed significant positive deviations in the left caudate nuclei (*t-value* = 3.31, *q*_*FDR*_ = 0.01), right caudate nuclei (*t-value* = 2.73, *q*_*FDR*_ = 0.04), left hippocampus (*t-value* = 3.28, *q*_*FDR*_ = 0.02), right hippocampus (*t-value* = 2.94, *q*_*FDR*_ = 0.03), left putamen (*t-value* = 4.17, *q*_*FDR*_ = 0.004), right putamen (*t-value* = 3.68, *q*_*FDR*_ = 0.007), left pallidum (*t-value* = 3.91, *q*_*FDR*_ = 0.005), and right amygdala (*t-value* = 3.01, *q*_*FDR*_ = 0.027), as well as the left lateral ventricle (*t-value* = 5.47, *q*_*FDR*_ = 0.0002) and right lateral ventricle (*t-value* = 4.40, *q*_*FDR*_ = 0.002).

**Figure 3.**
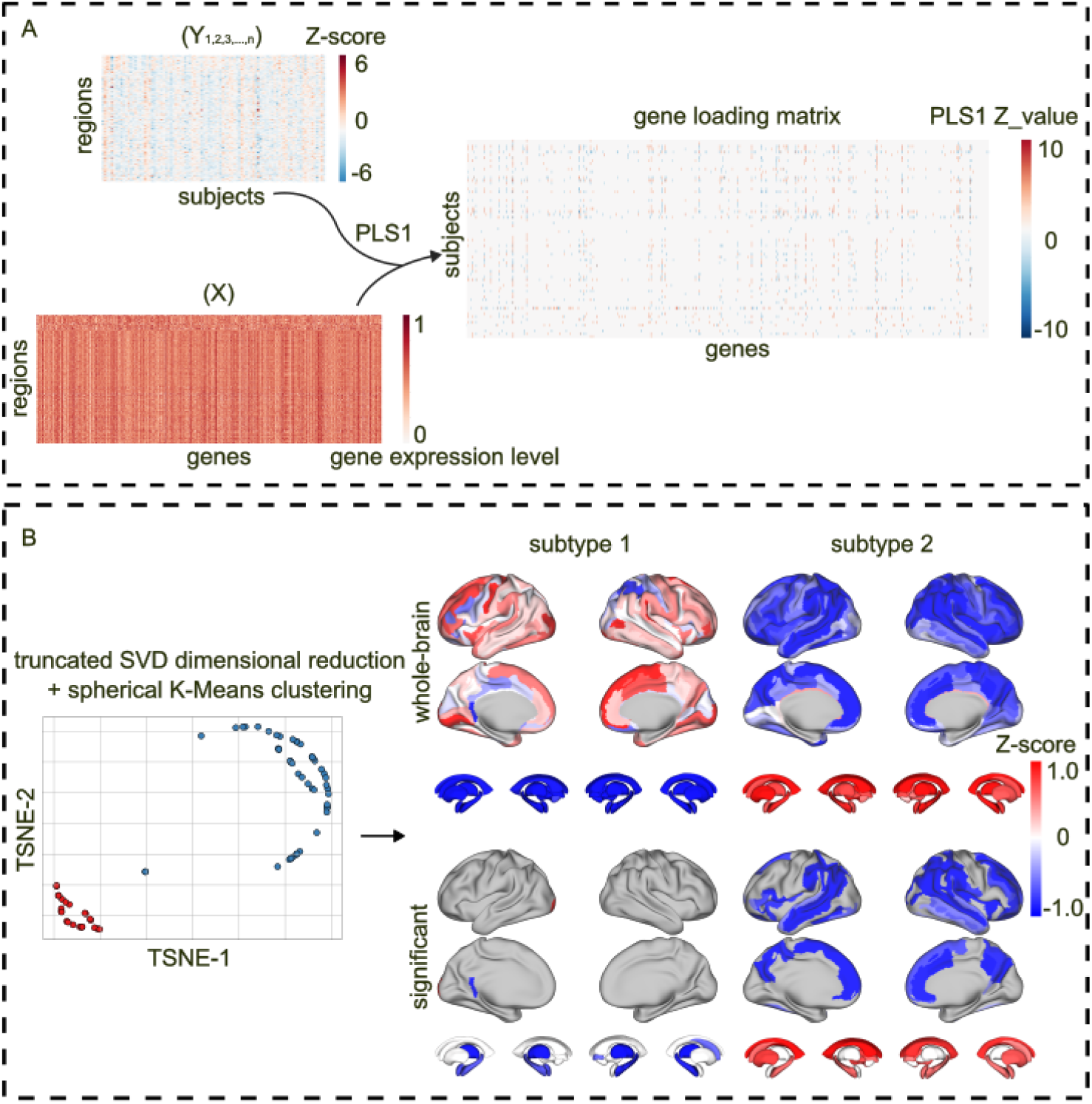
Transcriptomics-based EOS subtypes. **(A)** Schematic representation of the workflow combining EOS deviation maps with whole-brain gene expression from Allen Human Brain atlas to identify individual-level deviation-related transcriptomics. The subject × gene matrix (right) is computed using PLS analysis to reveal correlations between brain region-specific deviations and gene expression profiles at individual level. **(B)** (Left) t-SNE visualization of the data following dimensionality reduction using truncated SVD and clustering with spherical K-means, revealing two distinct subtypes of EOS patients. (Right, top) Brain-wide *Z-score*s for each subtype (subtype 1 and subtype 2) show the distribution of positive (red) and negative (blue) deviations, with the FDR-corrected deviations highlighted in the lower row compared with TD (P_FDR_ < 0.05). These results illustrate subtype-specific patterns of brain structural deviations in EOS. EOS, early-onset schizophrenia; TD, typically developing; PLS, partial least squares; PLS1, first PLS component; truncated SVD, truncated singular value decomposition; t-SNE, t-distributed stochastic neighbor embedding; FDR, false discovery rate.

### Clinical and biological relationships

To investigate clinical relevance of EOS subtypes, we further performed disease enrichment analyses and clinical correlations. For both subtypes, schizophrenia was the most significantly enriched disease factor (subtype 1 p = 1.85 × 10^-8^, subtype 2 p = 4.68 × 10^-11^). Additionally, both subtypes were also significantly enriched in gene sets associated with neurodevelopmental disorders, bipolar disorder, Alzheimer’s disease, and insomnia which were top-ranked by the value of -log_10_(pvalue), suggesting a shared genetic susceptibility between the two subtypes at the level of disease-related genes (**Figure 4A**). Regarding clinical correlations, we found no significant differences between the dimension scores of the PANSS of the subtypes (q_FDR_ < 0.05).

**Figure 4.**
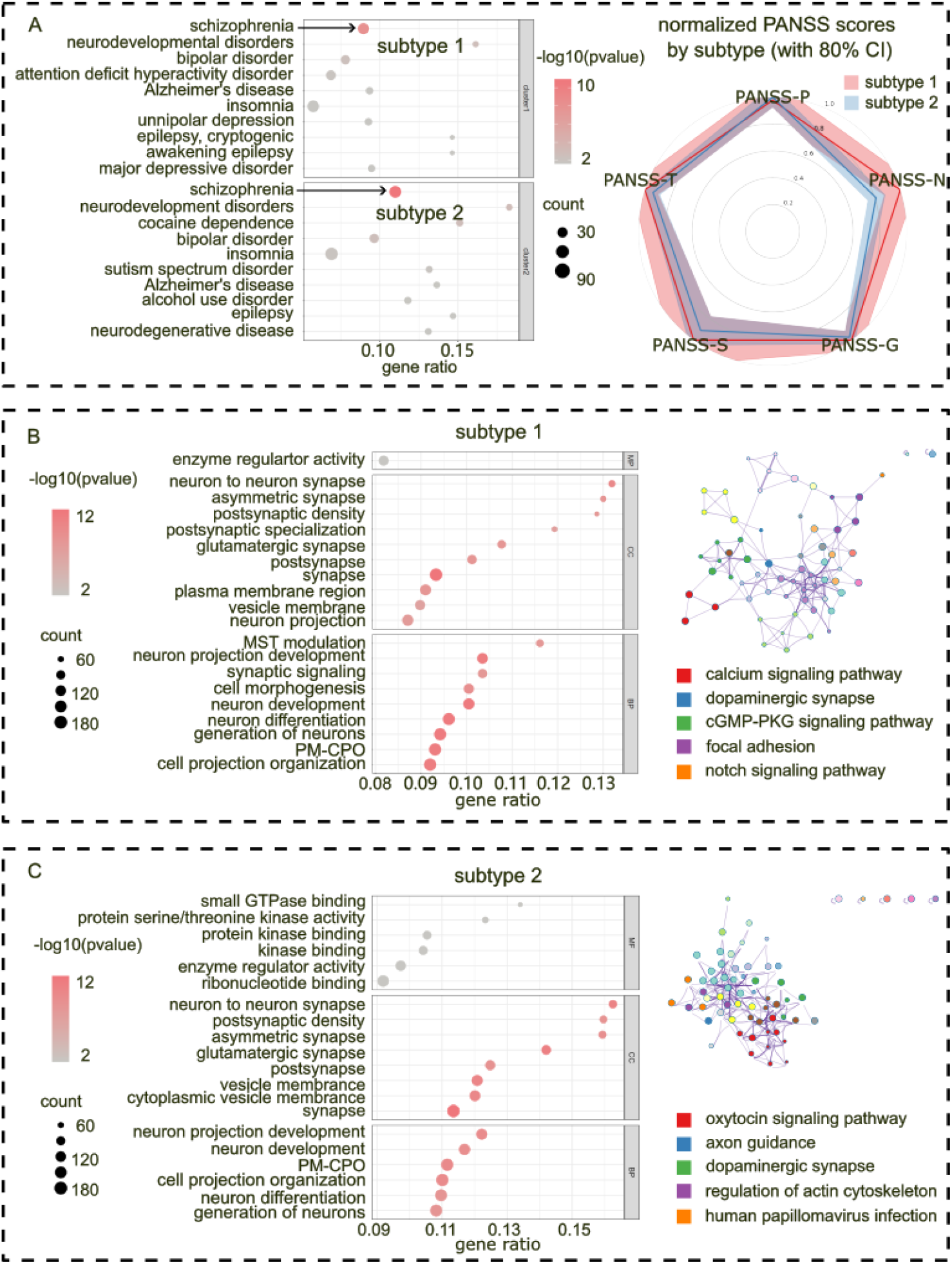
Clinical and gene enrichment analysis. **(A)** (Left) The gene enrichment analysis for each subtype illustrates the distribution of schizophrenia-related and other psychiatric disorder-related genes across the two subtypes. Disease factors were top-ranked by the value of -log_10_(p-value). (Right) The plot highlights the normalized PANSS scores (P, S, N, G) for each subtype, with 80% confidence intervals indicated. **(B & C)** GO Enrichment and KEGG Pathway Analysis for Each Subtype: On the left, the GO enrichment analysis highlights the key MF, CC, and BP significantly associated with each subtype. The right panel displays the KEGG pathway analysis for each subtype, listing the top five significantly enriched pathways. EOS, early-onset schizophrenia; TD, typically developing; count, the number of genes associated with each entry for the respective subtype; gene ratio, the proportion of genes related to that entry relative to the total number of genes in the pathway; PANSS, Positive and Negative Syndrome Scale; CI, Confidence Interval; GO, gene ontology; KEGG, Kyoto Encyclopedia of Genes and Genomes; MF, molecular function; CC, cellular component; BP, biological process; MST modulation, modulation of chemical synaptic transmission; PM-CPO, plasma membrane bounded cell projection organization.

To further investigate the physiological relevance of EOS subtypes, we performed GO enrichment analysis. We found distinct molecular functions and biological processes between the two subtypes. Specifically, subtype 1 was mainly associated with the molecular function of enzyme regulator activity (gene ratio = 0.0818, count = 116; Figure 4B), whereas subtype 2 was enriched in categories involving small GTPase binding (gene ratio = 0.1340, count = 43), protein serine/threonine kinase activity (gene ratio = 0.1233, count = 55), protein kinase binding (gene ratio = 0.1054, count = 92) and kinase binding (gene ratio = 0.1042, count = 101) (Figure 4C). For biological processes, subtype 1 was enriched in processes related to synaptic transmission, neuronal development, and differentiation, while subtype 2 was more strongly associated with neuronal plasticity and cellular structural organization. At the cellular component level, both subtypes were significantly enriched in neuronal structures such as synapses and postsynaptic densities, suggesting a common involvement in synaptic functionality (See **Table S4 & S5** for detailed statistical values).

To further explore the biological pathways underlying the differences between the two subtypes, we conducted the KEGG pathway analysis and found distinct enrichment patterns between the two subtypes. Subtype 1 was mainly enriched in the calcium signaling pathway, dopaminergic synapse, cGMP–PKG signaling pathway, and Notch signaling pathway, whereas subtype 2 showed significant enrichment in the oxytocin signaling pathway, axon guidance, dopaminergic synapse, regulation of the actin cytoskeleton, and human papillomavirus infection pathways. These results indicated that, although both subtypes shared synaptic-related molecular features, they diverged in the specific signaling and neurobiological pathways underlying neuronal regulation and plasticity.

## Discussion

In this study, we investigated brain structural heterogeneity of EOS by combing normative modeling with transcriptomic profiles. We identified individual-specific deviations across EOS patients, characterized by heterogeneous deviation effects in the orbital part of the left inferior frontal gyrus. We observed two neurobiologically distinct EOS subtypes, each showing unique spatial patterns of anatomical deviations. Despite both subtypes being enriched for neurodevelopmental disorders, they had divergent biological mechanisms.

Group-level comparisons suggested cortical thinning and enlarged subcortical nuclei in EOS, consistent with previous studies ^50–52^. In particular, we found negative deviations in the fronto-insular and temporo-parietal association cortices, and positive deviations in the left lateral ventricle. However, substantial inter-individual variability across EOS patients was found in brain structural deviations. The most heterogenous deviations were found in the lingual part of the left medial occipito-temporal gyrus, though it is noteworthy that the standard deviation of Z-scores across the whole brain was generally larger in EOS patients compared to healthy controls, indicating widespread heterogeneity—even if only the lingual region survived strict multiple comparisons correction. These results support the concept that EOS is clinical and biological heterogeneous and could not be adequately captured by a single uniform anatomical signature which may be obscured by the group-averaged analysis^23,53^.

EOS patients showed consistent abnormalities in the association cortical systems, which are involved in higher-order cognitive control, emotion processing, and language functions^54–56^, rather than reflecting uniform cortical shifts. This highlights the specificity of structural abnormalities in EOS, focusing on regions associated with complex neurocognitive processes. In contrast, positive deviations were relatively limited and largely confined to occipital/visual regions, indicating that regional increases were not the predominant feature of the group-level deviation profile. Within subcortical structures, enlargement of the bilateral lateral ventricles emerged as the most prominent FDR-significant difference, consistent with prior findings in schizophrenia-spectrum MRI studies^4,57^. Taken together, these results suggest that EOS deviations are highly variable at the individual level yet show partial convergence at the group level, motivating subsequent analyses aimed at identifying more neurobiologically coherent subgroups based on deviation profiles.

Two distinct subgroups with relatively stable separation in the molecular features space^47,49^ were identified using transcriptomic variability^9^. Subtype 1 showed a more focal cortical signature, with significant thickening in the left posterior–ventral cingulate gyrus and thinning in the left occipital pole, alongside significant volume reductions in key subcortical structures (bilateral hippocampi and thalamus, left amygdala, and right caudate). In contrast, subtype 2 exhibited more extensive cortical negative deviations across regions involved in higher-order cognitive functions, including areas related to emotion processing, sensory integration, and executive control. This subtype was simultaneously characterized by significant enlargement across multiple subcortical nuclei (bilateral caudate nuclei, hippocampi, and putamen; left pallidum; right amygdala) and bilateral lateral ventricular expansion. These subtype-specific deviation profiles help reconcile the combination of marked individual heterogeneity and partial group-level convergence observed in the case–control analyses. In fact, the stark differences in subcortical nuclei between the two subtypes may, to some extent, help explain the previously reported inconsistent findings in schizophrenia research, where both volume increases and decreases in subcortical structures have been observed^16,58,59^. In particular, subtype 2 largely conforms to the broader EOS pattern of widespread association-cortical negative deviations, whereas subtype 1 departs from this pattern by showing a comparatively restricted cortical signal together with prominent reductions in limbic–thalamic and striatal structures. This helps explain why previous group-level studies on cortical thinning in schizophrenia may have obscured subtype-specific characteristics, as the number of participants in subtype 2 is twice that of subtype 1, and the group-level deviation profile of subtype 2 is more similar to that of the entire EOS cohort^15,50,60^. Only a few regions showed significant differences, suggesting that the most distinct subtype differences are localized to a limited set of cortical and subcortical regions, rather than being distributed globally across the entire brain. Taken together, these findings underscore the utility of stratifying EOS using molecularly informed features, as it reveals distinct deviation profiles that might not be immediately apparent from case–control contrasts alone^1,17^.

The disease enrichment analysis revealed that both EOS subtypes were significantly enriched for schizophrenia-related gene sets, as well as for genes associated with neurodevelopmental disorders, bipolar disorder, Alzheimer’s disease, and insomnia. Recent studies have highlighted shared genetic risk factors between schizophrenia and various neurodevelopmental and psychiatric disorders^61,62^. This suggests that, despite differences in structural and molecular characteristics, the two subtypes share a common genetic susceptibility profile that links early-onset schizophrenia with other neurodevelopmental and psychiatric disorders. Interestingly, no significant differences were observed between the subtypes in the PANSS dimensions, indicating that these neurobiological distinctions likely underlie the same clinical phenotype. This reflects neurobiological heterogeneity within the disorder rather than distinct clinical categories. Previous studies have supported the “synaptic hypothesis of schizophrenia” which suggests that defects in synaptic pruning during adolescence, including both excessive pruning and insufficient pruning, may contribute to the etiology of schizophrenia^6,59,63^. In line with these findings, the GO enrichment analysis revealed that both EOS subtypes share common synaptic dysfunction as a core feature, with significant enrichment in synaptic and postsynaptic regions at the cellular component level.

Despite this similarity, the subtypes exhibited distinct molecular profiles. Subtype 1 was primarily enriched in pathways related to enzyme regulation and glutamatergic synapse, suggesting that dysfunctions in neurotransmitter dynamics may contribute to its pathology^64–66^. In contrast, Subtype 2 showed significant enrichment in pathways associated with small GTPase binding and kinase function, all of which play crucial roles in neuronal remodeling and long-term synaptic plasticity^67,68^. These findings may indicate that subtype 2 involves alterations in neuronal connectivity and synaptic restructuring, which are essential for maintaining brain network integrity and neuroplasticity. The KEGG pathway analysis further differentiated the two subtypes. Subtype 1 was enriched in pathways related to calcium signaling, dopaminergic synapse regulation, and neurodevelopment, suggesting its potential association with synaptic plasticity and neurotransmitter signaling, as suggested in previous studies^69,70^. These pathways are crucial for maintaining synaptic function and neurodevelopment, supporting the idea that subtype 1 might be more affected by issues related to neurotransmitter signaling and synaptic transmission. On the other hand, subtype 2 was significantly enriched in pathways related to oxytocin signaling, axon guidance, and actin cytoskeleton regulation, suggesting a stronger association with mechanisms that regulate cytoskeletal dynamics and neuronal connectivity^71,72^. These processes are critical for establishing and maintaining functional neuronal circuits, and their dysregulation may underlie the more extensive neuronal connectivity changes observed in subtype 2.

This study has some limitations that should be considered when interpreting the finds. First, the sample size, while sufficient for initial discovery, remains relatively small. This may limit the statistical power to detect subtle neuroanatomical or clinical differences between the identified EOS subtypes, potentially explaining the lack of significant clinical distinctions in the present analysis. A larger, multi-center cohort would improve the generalizability of the results, allow for more robust validation of the subtypes, and enable the detection of more nuanced clinical-biological associations. Second, the gene expression data used for enrichment analysis were derived from the Allen Human Brain Atlas, which is based on post⍰mortem brains of neurotypical adults. While this provides a valuable spatial reference, it does not capture disease⍰specific or developmental changes in gene expression that may be present in EOS. Future studies would benefit from integrating transcriptomic data from adolescent or young⍰adult patient populations, or using indirect measures such as polygenic risk scores coupled with imaging, to better model illness⍰related molecular processes. Third, our biological interpretation relied on gene set enrichment analysis, which is correlational and hypothesis⍰generating in nature. While it helps prioritize plausible pathways, it does not establish causal or mechanistic links. These findings warrant validation through more direct molecular approaches—such as in vitro or animal model experiments—and by examining relevant protein expression or epigenetic markers in patient⍰derived samples.

## Conclusion

This study reveals inter-individual heterogeneity in EOS anatomical deviations and their transcriptomics basis. Transcriptomics-related clustering analyses suggest two EOS subtypes, with subtype 1 reflecting disruptions in neurotransmitter signaling and synaptic transmission, and subtype 2 reflecting impaired cytoskeletal dynamics and neuronal connectivity. The findings emphasize the importance of molecularly informed stratification in understanding the complexities of EOS.

## Acknowledgements

The authors declare that they have no conflict of interest. This work was supported by the National Natural Science Foundation of China (62403105, 62333003, 82121003, 62373079), the Sichuan Science and Technology Program (2026NSFSC1509), the China Postdoctoral Science Foundation (2023M740524), the Sichuan Province Innovative Talent Funding Project for Postdoctoral Fellows, the Medical Engineering Cooperation Funds from University of Electronic Science and Technology of China (ZYGX2021YGLH201), the Science and Technology Department of Sichuan Province (2025ZNSFSC0734), the Health Commission of Sichuan Province (24CXTD11), Health Commission of Chengdu (2024141), the Sichuan Preventive Medicine Association (SYYXHPT202420).

